# Validity and Interpretation of Two-Sample Mendelian Randomization with Binary Traits

**DOI:** 10.1101/2024.06.09.598150

**Authors:** Zixuan Wu, Jingshu Wang

## Abstract

**Background:** Two-sample Mendelian randomization (MR) is widely applied to binary exposures and outcomes. Yet standard MR models rely on linear effect assumptions that are difficult to interpret for binary traits. Although liability-based interpretations have been suggested, it remains unclear whether conventional summary-data MR is formally justified in this setting or what causal parameter it identifies.

**Methods:** We develop a liability-threshold framework in which binary traits arise from underlying continuous liabilities. We derive explicit relationships between genome-wide association study (GWAS) coefficients obtained from logistic or linear regression on binary traits and marginal genetic associations on the liability scale. Under small genetic effects, typical for complex traits, observed-scale GWAS coefficients are approximately proportional to liability-scale associations.

**Results:** This proportionality implies that standard two-sample MR methods remain statistically coherent for binary traits. MR applied to binary exposures or outcomes estimates a scaled causal effect between underlying liabilities rather than an effect on the observed binary scale. The scaling factor depends primarily on trait prevalence and is directly computable. Simulations and UK Biobank analyses confirm that, after rescaling, MR using binary traits recovers liability-scale causal effects consistent with analyses based on continuous traits.

**Conclusions:** We provide a formal statistical justification for summary-data MR with binary traits and clarify the causal parameter being estimated. These results support routine MR practice for binary exposures and outcomes while enabling coherent interpretation of effect sizes.

**Key Messages:**

- The interpretation of two-sample MR with binary exposures or outcomes is often unclear because GWAS analyses are performed on the observed binary scale.
- Under a liability threshold framework with small genetic effects, GWAS coefficients from logistic or linear regression on binary traits are approximately proportional to genetic associations on an underlying continuous liability scale.
- Consequently, conventional summary-data MR applied to binary or ordinal traits remains valid and estimates a scaled causal effect between liabilities, requiring no modification of existing methods.

## 1 Introduction

Mendelian randomization (MR) uses genetic variants as instrumental variables to investigate the causal effects of modifiable exposures on health, behavioral, and social outcomes [10]. With the rapid growth of genome-wide association studies (GWAS), two-sample MR based on summary statistics has become the dominant approach in applied research [14]. Many MR analyses focus on binary or categorical traits, including disease status, smoking initiation, medication use, and educational attainment. As such traits are central to epidemiologic research, uncertainty about their treatment in MR analyses has important practical implications.

Binary traits pose challenges for standard MR frameworks. Unlike continuous traits, binary outcomes do not naturally support linear causal effect models, and binary exposures often represent thresholded versions of underlying continuous processes. Genetic variants may therefore influence an individual’s underlying risk without changing the observed binary exposure status, potentially affecting the outcome through the same biological pathway while appearing to violate exclusion restriction on the binary scale [5, 15]. Moreover, GWAS of binary traits are typically conducted using logistic regression, often in case–control samples, producing odds-ratio summary statistics [6]. Most two-sample MR methods, however, assume linear relationships between genetic associations with the exposure and outcome and summarize effects using a single population-level parameter. When the exposure is binary, this homogeneity assumption is difficult to justify: individuals classified as exposed may differ widely in their underlying risk or severity, making it implausible that changing exposure status from 0 to 1 has the same effect for all individuals. Consequently, it is unclear how such a parameter corresponds to meaningful causal contrasts[5, 17, 19].

An alternative interpretation for binary exposures invokes monotonicity and local average treatment effects (LATEs) [1]. In MR studies using many genetic instruments, however, monotonicity implies instrument-specific complier subgroups, so different variants may identify different local effects. Combining them therefore requires strong homogeneity assumptions, further complicating interpretation [5, 19]. Standard two-sample MR applied to binary traits is thus often regarded as potentially problematic.

The liability threshold model, which treats binary traits as dichotomizations of underlying continuous variables, has long been used in genetic and epidemiologic research [11, 13]. Recent work in MR has suggested interpreting MR analyses of binary or ordinal traits on a liability scale [5, 15]. However, these discussions emphasize interpretational difficulty and additional assumptions. No formal statistical derivation has established whether conventional summary-data MR applied to binary traits is valid under a liability interpretation, or precisely what parameter it identifies.

In this paper, we provide a formal statistical justification for conventional two-sample MR with binary exposures and outcomes under the liability framework. We derive explicit relationships between GWAS coefficients from logistic or linear regression on binary traits and genetic associations on the underlying liability scale, quantify the corresponding prevalence-dependent scaling factors under cohort and case–control designs, and characterize approximation accuracy. Under small genetic effects, observed-scale GWAS coefficients are approximately proportional to liability-scale associations. Consequently, conventional two-sample MR applied to binary traits remains statistically coherent without modification: it estimates a scaled causal effect between underlying liabilities rather than an effect on the observed binary scale. This interpretation extends to more complex MR frameworks, including multivariable, within-family, and life-course MR [2, 4, 8, 18, 22], supporting principled use of binary and ordered categorical traits across epidemiologic applications.

## 2 Conceptual framework: MR on the liability scale

### 2.1 Observed binary traits and latent liabilities

Figure 1a shows the standard MR setting in which genetic variants, or single nucleotide polymorphisms (SNPs), are used as instruments for an observed exposure *X* to estimate its causal effect on an observed outcome *Y*, in the presence of unmeasured confounders *U*. Standard MR analyses rely on assumptions that genetic variants are independent of *U* and satisfy the exclusion restriction, meaning they affect the outcome only through the exposure. When both *X* and *Y* are continuous traits, these assumptions are imposed directly on the observed variables.

**Figure 1:**
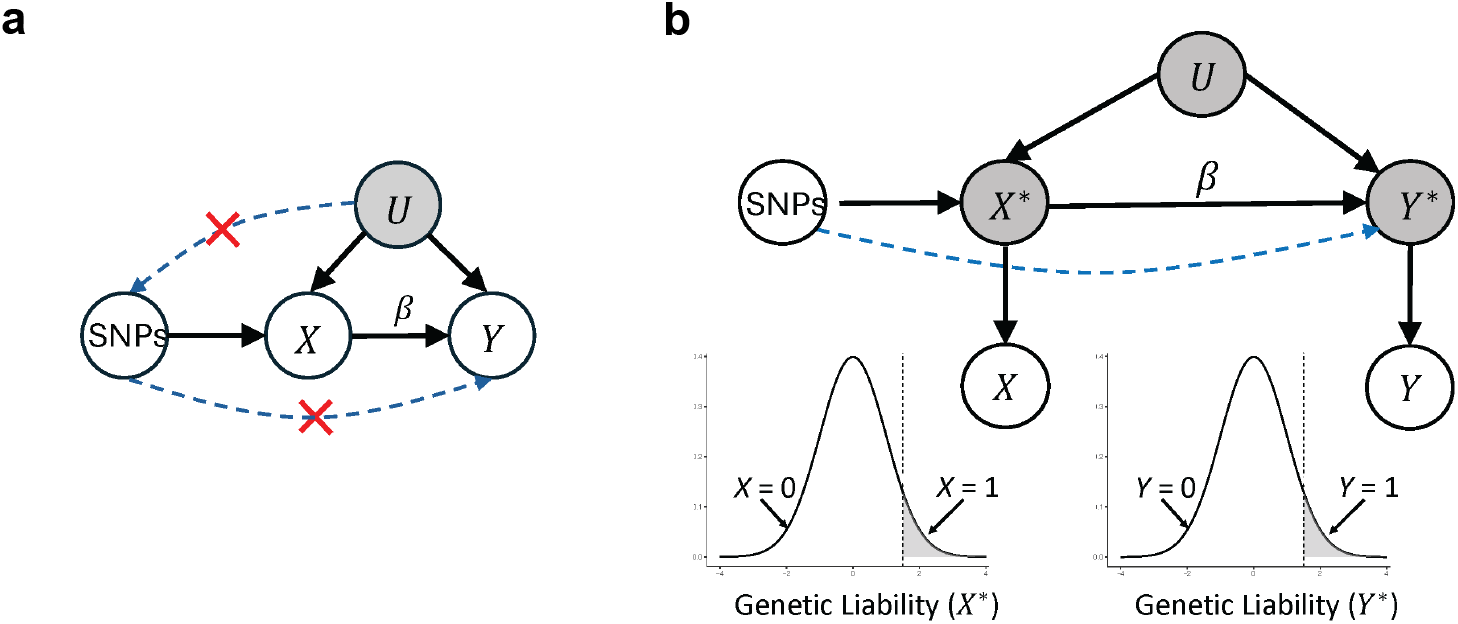
Conceptual framework for MR with binary traits. a) Classical MR framework in which genetic variants (SNPs) influence the exposure *X*, which in turn affects the outcome *Y*, while remaining independent of unmeasured confounders *U*. The red crosses denote the independence and exclusion restriction assumptions. b) Liability-scale interpretation. The observed traits *X* and *Y* may be binary and arise from underlying continuous liabilities *X*^*∗*^ and *Y* ^*∗*^. The causal effect *β* is defined between the liabilities, while the observed binary traits result from thresholding these latent variables. The dashed blue arrow indicates possible horizontal pleiotropy.

However, when exposures or outcomes are binary, it is often more realistic to view the observed phenotypes as coarse measurements of underlying continuous processes. For example, disease diagnoses typically reflect whether an individual’s underlying risk exceeds a clinical threshold, and behavioral or social exposures often reflect whether latent tendencies cross reporting or classification boundaries. Figure 1b illustrates this perspective, in which each observed binary trait arises from a latent continuous liability that summarizes an individual’s overall underlying risk for the trait.

Specifically, we assume that the observed exposure *X* equals one when an unobserved exposure liability *X*^⋆^ exceeds a trait-specific cutoff *t*_*X*_, that is, *X* = 1(*X*^⋆^ *> t*_*X*_), and similarly, the observed outcome *Y* equals one when an unobserved outcome liability *Y* ^⋆^ exceeds a cutoff *t*_*Y*_. This framework applies whenever either the exposure or outcome is binary; if a trait is continuous, it coincides with its own liability. Individuals with the same observed binary status may differ substantially in their underlying liabilities, and genetic variants influence these latent risks rather than directly determining observed status.

### 2.2 Causal effects and MR assumptions on the liability scale

Under the liability framework, causal relationships are defined between the continuous liabilities *X*^⋆^ and *Y* ^⋆^, rather than between the observed binary variables *X* and *Y*. The causal effect of interest is the effect of *X*^⋆^ on *Y* ^⋆^.

Accordingly, the core MR assumptions, including independence of genetic variants from confounders, exclusion restriction, and assumptions concerning horizontal pleiotropy, are imposed on the latent liabilities. That is, genetic variants are assumed to influence the outcome liability only through their effects on the exposure liability, except for any explicitly modeled pleiotropic pathways.

Defining causal effects on the liability scale avoids interpreting MR estimates as effects of crossing diagnostic thresholds and instead corresponds to effects of incremental changes in latent risk. Heterogeneity in observed binary responses is therefore compatible with a stable population-level causal effect between *X*^⋆^ and *Y* ^⋆^.

### 2.3 Implications for GWAS and summary-data MR

Although causal effects and MR assumptions are defined on the liability scale, both GWAS and MR analyses rely on observed data. In practice, GWAS of binary traits estimate associations between genetic variants and observed phenotypes using logistic or linear regression, producing summary statistics on the odds-ratio or linear probability scale. A key question is whether these observed-scale associations preserve information about genetic effects on the liabilities.

The key insight is that, because individual genetic variants typically have small effects on complex traits, the change in the probability of a binary outcome induced by a single variant can be approximated by a local linear shift in the corresponding liability. Consequently, GWAS summary statistics for binary traits are approximately proportional to the corresponding marginal genetic associations on the liability scale, with proportionality determined by trait prevalence and study design. Standard two-sample MR applied to these summary statistics therefore identifies a scaled version of the causal effect between liabilities.

In the next section, we formalize this relationship and derive the corresponding scaling factors for summary statistics obtained from logistic and linear GWAS under both cohort and case-control sampling.

## 3 Liability-Scale interpretation of GWAS summary statistics

### 3.1 Genetic associations on the liability scale

When a trait is observed on a continuous scale, GWAS summary statistics are conventionally obtained from linear regression. Let *X*^⋆^ denote the exposure liability and *Z*_*j*_ denote SNP *j*. The marginal genetic association is defined through the regression

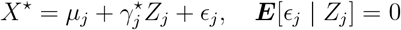

The coefficient 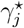 represents the marginal association between SNP *j*. Analogous notation 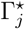 is used for the outcome liability *Y* ^⋆^. When only a binary trait *X* is observed, the liability *X*^⋆^ is unobserved, and 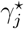 cannot be estimated directly.

### 3.2 GWAS coefficients for binary traits under the liability model

We now quantify how GWAS summary statistics obtained from binary traits relate to genetic associations with the underlying liabilities. For analytical clarity, we adopt the conventional liability threshold model, assuming that the liability follows a standard normal distribution. Extensions to more general liability distributions are discussed in Supplementary Section S3.3.

Let 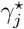 denote the marginal association between SNP *j* and the exposure liability *X*^⋆^, as defined in Section 3.1. In practice, GWAS summary statistics for a binary exposure are obtained by regressing the observed trait *X* on each SNP. Let 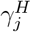 and 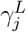 denote the expected regression coefficients from logistic and linear regression, respectively, when analyzing the binary exposure, with linear regression applied to the standardized binary outcome.

Under the liability threshold model and assuming that individual SNP effects on liability are small, GWAS regression coefficients from the observed binary trait are approximately proportional to the corresponding liability-scale genetic associations. Specifically, it can be shown (see Supplementary Section S1.1-S1.2) that

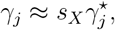

where *γ*_*j*_ denotes the GWAS regression coefficient for SNP *j* (from either logistic or linear regression) on *X* and *s*_*X*_ is a scaling factor that depends on the prevalence of the trait, the regression model, and the sampling design, but is constant across SNPs within a study.

Under cohort sampling, these scaling relationships take the form

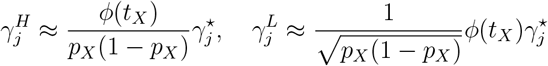

where *p*_*X*_ = P(*X* = 1) denotes the population prevalence of the binary exposure, *t*_*X*_ = Φ^−1^(1 − *p*_*X*_) is the corresponding liability cutoff, and *ϕ* denotes the standard normal density.

Under case–control sampling, logistic regression coefficients remain approximately unchanged, whereas linear regression coefficients depend additionally on the case fraction in the sample. In this setting, the expected coefficients satisfy

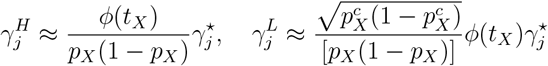

where 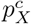 denotes the proportion of cases in the case–control sample. The difference between cohort and case–control designs reflects the fact that logistic regression is approximately invariant to retrospective sampling, whereas linear regression depends on the proportion of cases in the analyzed sample. Consequently, logistic regression coefficients follow the same scaling under both designs, while linear regression coefficients involve an additional factor determined by the case fraction.

Figure 2a displays the scaling factor *s*_*X*_ as a function of prevalence *p*_*X*_ for logistic and linear regression, respectively, under cohort sampling and standard normal liability. These curves describe how observed-scale GWAS coefficients can be rescaled to recover liability-scale effects. For logistic regression, the scaling factor is close to the commonly used logistic–probit conversion factor of 1.7 over a wide range of prevalences [7], but deviates substantially when prevalence is very low or very high. For linear regression, the scaling factor varies more strongly with prevalence and is generally smaller in magnitude.

**Figure 2:**
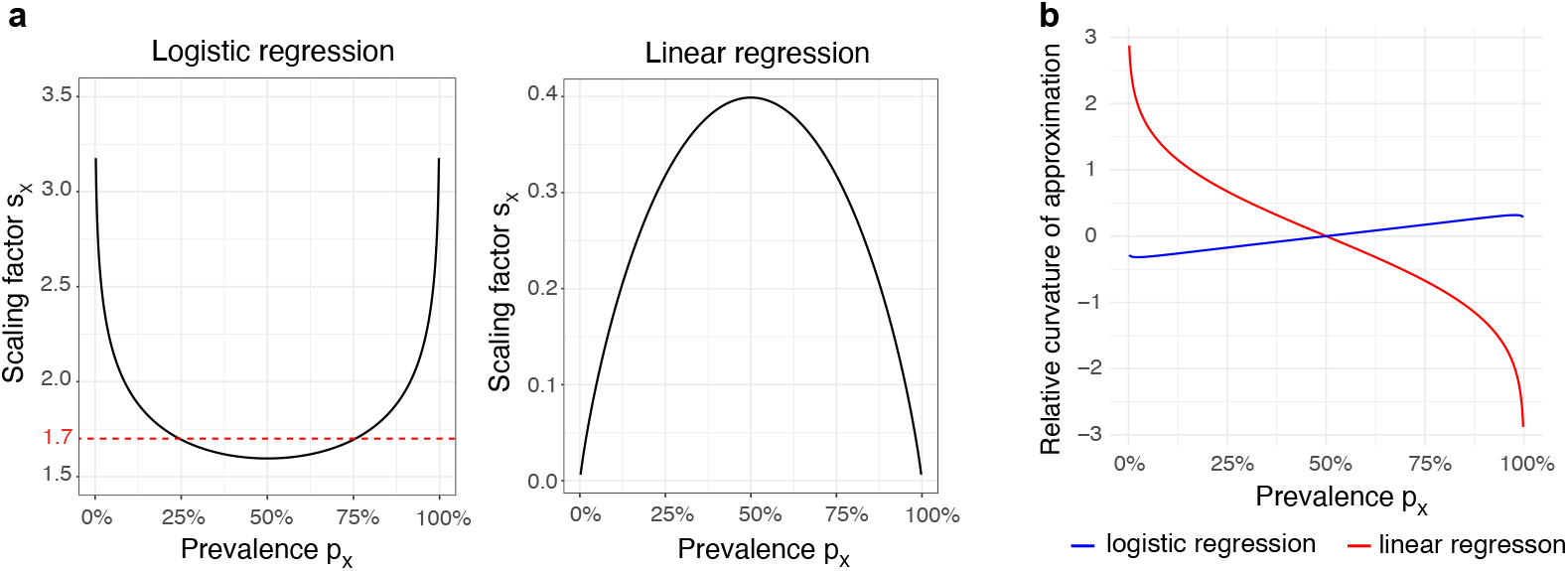
a): Scaling factor *s*_*X*_, the ratio between GWAS regression coefficients for a binary trait and liability-scale genetic effects, shown as a function of the prevalence *p*_*X*_ = P(*X* = 1) under the standard normal liability model and cohort sampling. The left panel corresponds to logistic regression and the right panel to linear regression (applied to the standardized binary trait). The dotted red line indicates the commonly used logistic–probit conversion factor of 1.7. b) Relative curvature of the approximation linking liability shifts to observed-scale regression coefficients, for logistic (blue) and linear (red) regression. Larger curvature magnitude indicates greater deviation from proportionality.

### 3.3 Approximation accuracy and prevalence dependence

The scaling relationships in Section 3.2 arise from first-order Taylor approximations linking small liability shifts to changes in binary trait probabilities and GWAS regression coefficients (Supplementary Section S1.1). The accuracy depends on higher-order terms in this expansion, governed by the local curvature of the probability function near the liability cutoff.

Intuitively, the relative curvature indicates the relative accuracy of linearizing the association between binary traits and the genetic variants under the liability model. Formal expressions and complete derivations are given in Supplementary Section S1.3. When prevalence is close to 0 or 1, the relative curvature deviates more from 0, indicating that the observed-scale regression coefficients may deviate more substantially from proportionality to liability-scale genetic effects (Figure 2b). By contrast, when prevalence is moderate, the relative curvature is closer to 0, and the proportional relationships are more accurate. Moreover, logistic regression exhibits substantially smaller curvature than linear regression when modeling binary traits, explaining why logistic GWAS coefficients remain closer to liability-scale associations across a wider range of prevalences.

### 3.4 Consequences for two-sample MR

Under standard MR assumptions imposed on the liabilities, genetic associations satisfy

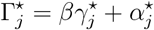

where *β* represents the causal effect of exposure liability on the outcome liability and 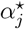 denotes the horizontal pleiotropic effects. Here *β* may represent the homogeneous causal effect, an average derivative effect under a smooth nonlinear model, or a summary effect under a more general structural equation linking *X*^⋆^ and *Y* ^⋆^ (See Supplementary Section S2 for details.)

When GWAS summary statistics are obtained from observed binary traits, Section 3.2 implies that the expected observed-scale coefficients satisfy 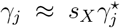 and 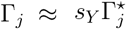, where Γ_*j*_ denotes the GWAS regression coefficient for SNP *j* on the outcome *Y*.

Substituting into the MR model yields

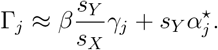

Thus, standard two-sample MR applied to observed binary traits estimates a scaled version of the liability-scale causal effect, with scaling factor *s*_*Y*_ */s*_*X*_, determined by prevalence, regression model, and sampling design, which are typically known or can be estimated in practice. Importantly, no modification of standard MR procedures is required; only the interpretation of the estimated effect changes.

In practice, GWAS analyses typically adjust for additional covariates such as age, sex, and ancestry principal components. Under standard MR assumptions, genetic variants are independent of these covariates, and the proportionality between observed-scale GWAS coefficients and liability-scale genetic effects continues to hold approximately after covariate adjustment. For linear regression, independence between genetic variants and covariates implies that adjustment does not change the marginal SNP coefficient. For logistic regression, although non-collapsibility alters odds ratios, proportionality is preserved under small genetic effects. Formal results are provided in the Supplementary Section S3.1.

The same reasoning extends naturally beyond binary traits. Ordered categorical traits can be viewed as discretizations of an underlying continuous liability using multiple thresholds. Under such models, GWAS coefficients from proportional odds or related regression models are likewise proportional to liability-scale genetic associations, with prevalence-dependent scaling factors (see Supplementary Section S3.2 for formal derivations).

## 4 Empirical evaluations

### 4.1 Simulation study: validating scaling of GWAS and MR estimates

We conduct simulations to evaluate whether GWAS coefficients from binary traits follow the predicted proportionality to liability-scale genetic effects and whether MR recovers the liability-scale causal parameter after rescaling.

We generate a latent continuous exposure *X*^⋆^ from a linear model involving 100 independent SNPs (Supplementary Section S4). Minor allele frequencies are sampled from the empirical UK Biobank distribution, and SNP effects are drawn from a standard normal distribution. Gaussian noise is added so that the SNPs jointly explain approximately 60% of the variance in *X*^⋆^, which is then standardized.

Binary traits are created by dichotomizing *X*^⋆^ at thresholds corresponding to prevalences of 50%, 20%, 5%, 1%, and 0.1%. These settings represent common, moderately common, rare, and very rare traits. GWAS coefficients are obtained using logistic and linear regression on the binary trait and compared to linear regression coefficients from the continuous liability. Reported values are medians across 100 simulation replicates.

As shown in Figure 3a, binary-trait GWAS coefficients are approximately proportional to the corresponding liability-scale effects. Dividing by the theoretical scaling factor *s*_*X*_ aligns the estimates closely with the identity line. Accuracy is highest for logistic regression and moderate prevalences. Additional prevalences are shown in Supplementary Figure S1a. In addition, Supplementary Figure S1b shows that simulated effect sizes fall within the range observed in empirical GWAS, confirming that the simulation operates in a realistic effect-size regime.

**Figure 3:**
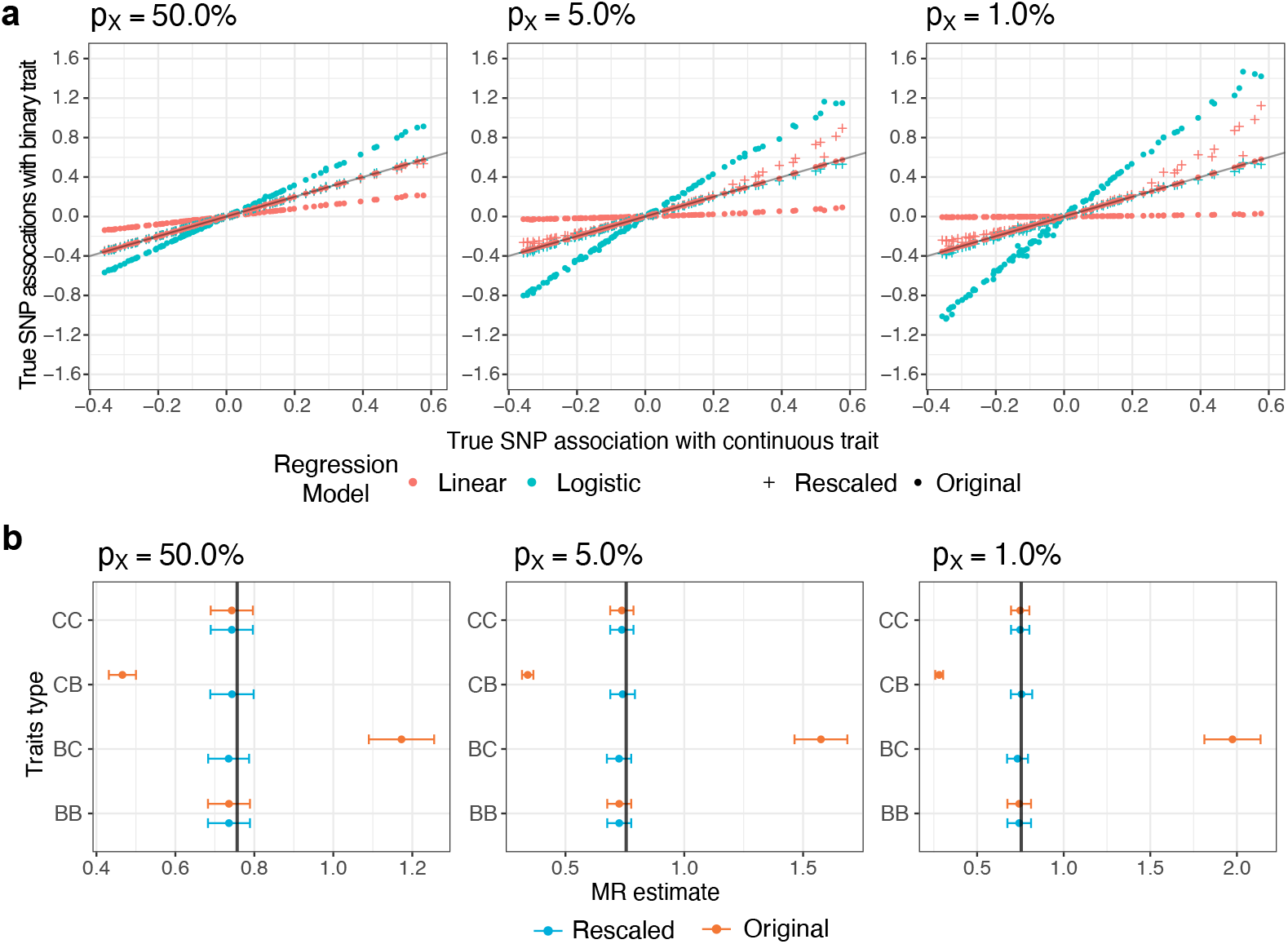
a) True per-SNP associations on the underlying continuous trait versus associations estimated from the observed binary trait using logistic or linear regression, shown before and after rescaling by the factor *s*_*X*_, for prevalences *p*_*X*_ = 50%, 5% and 1%. Rescaled estimates are obtained by dividing the binary-trait regression coefficients by *s*_*X*_. Points represent median estimates across 100 simulation replicates. The black line indicates *y* = *x*. b) MR 95% confidence intervals computed using GRAPPLE under moderate pleiotropy, shown before and after rescaling, for four exposure-outcome combinations: continuous-continuous (CC), continuous-binary (CB), binary-continuous (BC), and binary-binary (BB). When traits are binary, exposure and outcome share the same prevalence. Results are shown for one representative simulation replicate. The true causal effect is approximately 0.75 (black vertical line).

We next evaluate MR estimation under two pleiotropy settings: small and moderate horizontal pleiotropy. For each, we consider four exposure–outcome combinations: both traits continuous, exposure continuous with binary outcome, exposure binary with continuous outcome, and both traits binary. When binary traits were involved, MR estimates target the scaled parameter *βs*_*Y*_ */s*_*X*_, and we therefore compare both observed and rescaled estimates. MR is performed using GRAPPLE [20], which provides robust inference in the presence of weak instruments and random horizontal pleiotropy.

Under moderate pleiotropy (Figure 3b), unscaled estimates can vary substantially across trait definitions, whereas rescaled estimates closely match the continuous–continuous benchmark. Under small pleiotropy (Supplementary Figure S1c), the pattern holds, with slightly greater precision for continuous traits due to information loss from dichotomization. Across scenarios, appropriate scaling restores agreement with the liability-scale effect.

Overall, differences between continuous and binary MR analyses arise primarily from scale transformations; prevalence-based correction recovers the underlying causal parameter.

### 4.2 Application to UK Biobank

We further illustrate these findings using UK Biobank data, a large prospective cohort of more than 500,000 individuals recruited between the ages of 40 and 69 years. We consider body mass index (BMI) as the exposure and systolic blood pressure (SBP) as the outcome. Binary versions of are defined using standard clinical cutoffs: obesity (BMI *>* 30; prevalence 23.8%) and hypertension (SBP *>* 140 mmHg; prevalence 46.7%).

Independent genetic instruments for BMI are selected via LD clumping using SNPs associated at p-values *<* 10^−4^ in GIANT [16, 21]. Marginal associations are estimated using linear regression for continuous traits and logistic regression for binary traits, without additional covariate adjustment. MR is performed using IVW [3] and GRAPPLE under four exposure–outcome definitions: BMI to SBP, BMI to hypertension, obesity to SBP, and obesity to hypertension. When binary traits are involved, estimates are also rescaled to the liability scale.

Figure 4 shows that observed-scale estimates differ across trait definitions, whereas rescaled estimates converge to nearly identical confidence intervals. Apparent discrepancies between continuous and binary MR analyses largely reflect scale differences rather than substantive changes in causal inference. After correction, binary-trait MR estimates are directly comparable to continuous-trait results.

**Figure 4:**
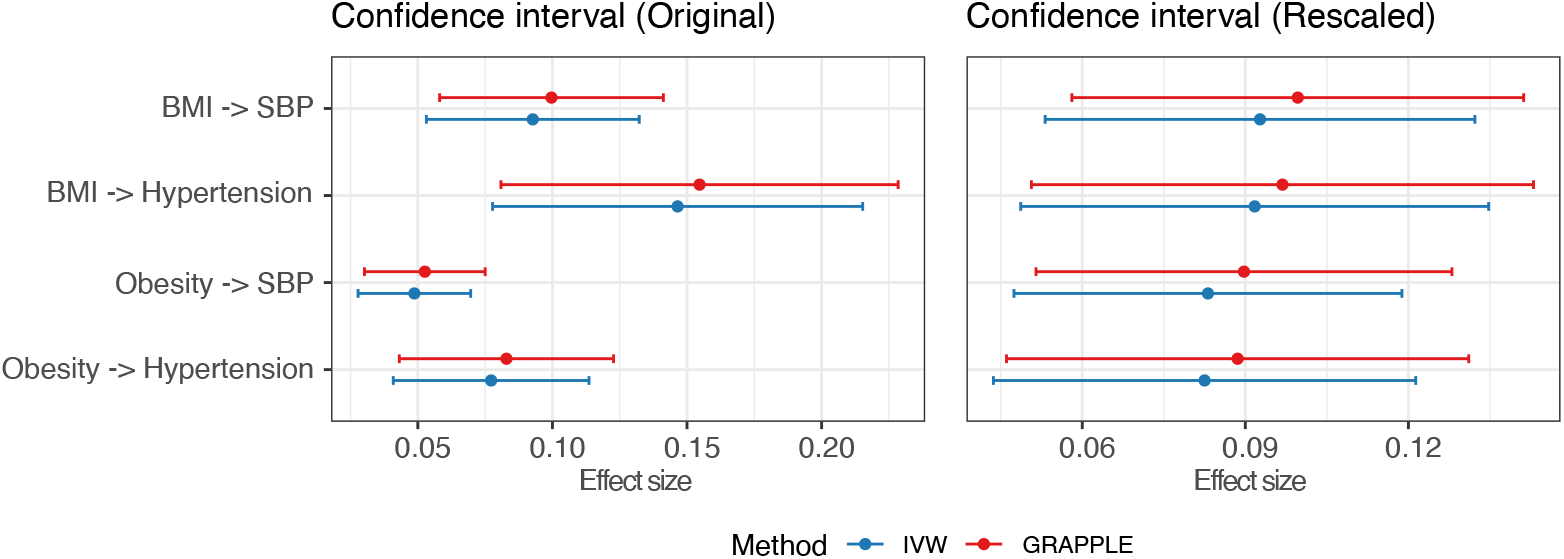
MR 95% confidence intervals for the causal effect of BMI on SBP under different exposure–outcome definitions, estimated using IVW and GRAPPLE. The left panel displays estimates obtained from observed-scale summary statistics, while the right panel shows results after rescaling binary-trait associations to the liability scale. After rescaling, estimates across continuous and binary trait definitions align closely and yield consistent conclusions.

## 5 Discussion

In this paper, we provide a formal justification for applying conventional two-sample MR methods to binary exposures and outcomes under a liability framework. When genetic effects are small, GWAS summary statistics from logistic or linear regression on binary traits are approximately proportional to genetic associations on an underlying continuous liability scale. Standard summary-data MR estimators therefore remain valid without modification: they identify a scaled causal effect between exposure and outcome liabilities, with scaling determined by prevalence and study design. Because this interpretation relies only on the proportionality between observed-scale and liability-scale associations, it supports not only univariable MR but also more complex settings, including multivariable, within-family, and life-course MR, when traits are binary.

These results clarify a longstanding concern in applied MR. Binary traits are often viewed as requiring alternative estimators or stronger assumptions. We show that, under the liability threshold model and small genetic effects typical of GWAS, conventional summary-data MR already recovers a well-defined causal parameter on the liability scale. Differences between continuous and binary analyses largely reflect scale transformations rather than violations of instrumental variable assumptions.

The derivations extend earlier theoretical work on binary outcomes [23] by providing a unified framework for both binary exposures and outcomes explicitly and by characterizing how scaling depends on prevalence and sampling design. Methods specifically tailored to binary outcomes [9, 12] often require individual-level data, whereas our approach applies directly to widely available summary statistics.

Our interpretation still has several limitations. We assume that binary traits arise from a common threshold applied to a continuous underlying risk. In practice, diagnostic or behavioral thresholds may vary across subgroups. Our proportionality results rely on small SNP effects and may be less accurate for variants with large effects or extremely rare traits. Additionally, although proportionality is preserved under covariate adjustment and ordinal traits, extreme prevalences may reduce approximation accuracy.

Overall, binary exposures and outcomes need not be viewed as obstacles to two-sample MR. Rather, they can be interpreted as discretizations of continuous liabilities, with standard summary-data MR recovering scaled causal effects between these latent traits. This perspective clarifies interpretation and strengthens the practical foundations of MR in epidemiologic research.

## Ethics approval

The UK Biobank study is approved by the National Health Service’s Health Research Authority (ref. 21/NW/0157; UK Biobank Application Number 27386).

## Acknowledgments

We thank the University of Chicago Research Computing Center for computational support, and we are grateful to Xin He for helpful comments.

## Author contributions

J.W. conceived and supervised the study. J.W. and Z.W. developed the framework. Z.W. implemented the proposed method, conducted the simulation studies, and performed the real data analyses. J.W. and Z.W. wrote the manuscript.

## Data availability

Due to the terms of the original participant consent and UK Biobank data access policies, individual-level data used in this study cannot be made publicly available in the manuscript, supplementary materials, or a public repository. However, the data are available to qualified researchers upon application to the UK Biobank Access Management Team. Investigators seeking access must submit a research proposal, which will be reviewed and approved by UK Biobank. Further information regarding the application process is available from UK Biobank. The phenotypes analyzed in this study include body mass index (BMI; field 21001) and systolic blood pressure (SBP; field 4080).

