## Supplemental Materials for "Validity and Interpretation of Two-Sample Mendelian Randomization with Binary Traits"

### Supplementary Materials

These Supplementary Materials are organized as follows. Section S1 provides detailed mathematical derivations of the proportionality and approximation results. Section S2 discusses general structural models for causal effects on the liability scale. Section S3 presents extensions, including covariate adjustment, ordered traits, and general liability distributions. Section S4 describes simulation settings and implementation details. Section S5 provides additional figures.

#### S1 Derivation and Accuracy of the Proportionality Approximation

##### S1.1 Derivation of the observed–liability proportionality

We begin by recalling the definition of marginal SNP effects when the underlying trait is continuous. When $X^\star$ is directly observed, the marginal association between a SNP $Z_j$ and $X^\star$ is defined via the linear projection

$$X^\star = \mu_j + \gamma_j^\star Z_j + \epsilon_j, \qquad \boldsymbol{E}[\epsilon_j \mid Z_j] = 0.$$

Throughout, we assume that the latent trait $X^\star$ follows a standard normal distribution with cumulative distribution function (CDF) $\Phi$. (See Section S3.3 for general distributions.) Consistent with the highly polygenic architecture of complex traits, we further assume that the effect of any single SNP is small, i.e., $\gamma_j^\star = o(1)$. Under this assumption, the contribution of $\mu_j$ and $\epsilon_j$ dominates, so that $\mu_j + \epsilon_j \approx X^\star$.

We consider a binary trait $X$ generated by thresholding the latent variable $X^\star$ at a cutoff $t_X$. Under this liability model,

$$\begin{aligned} \mathrm{P}(X=1) &= \mathrm{P}(X^\star \geq t_X) \\ &= \mathrm{P}(-\mu_j - \epsilon_j \leq \gamma_j^\star Z_j - t_X) \\ &\approx \Phi(\gamma_j^\star Z_j - t_X). \end{aligned} \tag{1}$$

Now suppose we fit a generalized linear model for the binary outcome $X$ using a link function $g(\cdot)$. Then

$$\begin{aligned} g(\mathrm{P}(X=1)) &\approx g\big(\Phi(\gamma_j^\star Z_j - t_X)\big) \\ &\approx (g\Phi)(-t_X) + (g\Phi)'(-t_X)\, \gamma_j^\star Z_j, \end{aligned} \tag{2}$$

where the second step is based on the first-order Taylor expansion of the function $g\Phi(\cdot)$ at $-t_X$. The above approximations show that $\gamma_j$ obtained from the binary-trait GWAS is approximately proportional to the marginal effect $\gamma_j^\star$ on the latent continuous scale:

$$\gamma_j \approx s_X \gamma_j^\star, \tag{3}$$

where the scaling factor $s_X = (g\Phi)'(-t_X)$ does not depend on the SNP index $j$.

Finally, recall that by definition $\mathrm{P}(X=1) = \mathrm{P}(X^\star > t_X) = \Phi(-t_X)$. Letting $p_X = \mathrm{P}(X =$

1) denote the prevalence of the binary trait, we can rewrite the scaling factor as

$$s_X = (g\Phi)'\big(\Phi^{-1}(p_X)\big)\,, \tag{4}$$

making explicit that $s_X$ depends only on the chosen link function $g(\cdot)$ and the trait prevalence.

#### S1.2 Explicit scaling factors under different models and designs

We now calculate the explicit forms of the scaling factor $s_X$ for two commonly used GWAS models for binary traits: logistic regression and linear regression. Throughout this section, we continue to assume a standard normal liability, $X^\star \sim N(0,1)$.

**Logistic regression.** Let $H(x) = \exp(x)\{1+\exp(x)\}^{-1}$ denote the logistic function. Logistic regression fits the model

$$H^{-1}(\mathrm{P}(X=1)) = \gamma_{0j} + \gamma_j^H Z_j.$$

Combining (3) and (4), the resulting regression coefficient satisfies

$$\gamma_j^H \approx s_X \gamma_j^\star, \qquad s_X = (H^{-1}\Phi)'\big(\Phi^{-1}(p_X)\big) = \frac{\phi(t_X)}{p_X(1-p_X)}, \tag{5}$$

where $\phi(\cdot)$ and $\Phi(\cdot)$ denote the standard normal density and cumulative distribution function, respectively.

**Linear regression.** In large-scale biobank studies [ukb], it is also common to apply linear regression directly to binary traits. Although this approach does not explicitly model the binary nature of the outcome, the resulting summary statistics remain approximately proportional to the latent effect $\gamma_j^\star$ under the small-effect assumption, albeit with reduced accuracy.

Specifically, linear regression fits

$$\mathrm{P}(X=1) = \mu_{0j} + \gamma_j^L Z_j,$$

corresponding to the identity link. Applying the same approximation as above yields

$$\gamma_j^L \approx s_X \gamma_j^\star, \qquad s_X = \Phi'\big(\Phi^{-1}(p_X)\big) = \phi(t_X).$$

In practice, the binary trait is often standardized by its standard deviation $\sqrt{p_X(1-p_X)}$ prior to regression. After standardization, the estimated coefficient satisfies

$$\gamma_j^L \approx \frac{1}{\sqrt{p_X(1-p_X)}}\, s_X \gamma_j^\star = \frac{\phi(t_X)}{\sqrt{p_X(1-p_X)}}\, \gamma_j^\star. \tag{6}$$

**Case–control sampling.** Under case–control sampling, logistic regression coefficients are invariant to the sampling scheme because they correspond to log odds ratios. Let $\gamma_j^{H,c}$ denote the logistic regression coefficient estimated from a case–control study. Consequently,

$$\gamma_j^{H,c} = \gamma_j^H \approx \frac{\phi(t_X)}{p_X(1-p_X)}\, \gamma_j^\star. \tag{7}$$

In contrast, linear regression coefficients depend explicitly on the case fraction in the sample. Assume first that the binary trait $X$ is not standardized in either the cohort or the case–control analysis, and let $\gamma_j^L$ and $\gamma_j^{L,c}$ denote the corresponding linear regression coefficients. Results from [5, 8] imply

$$\gamma_j^{L,c} \approx \frac{p_X^c(1-p_X^c)}{p_X(1-p_X)}\gamma_j^L,$$

where $p_X^c$ denotes the proportion of cases in the case–control study.

Now suppose that $X$ is standardized using the cohort prevalence $p_X$ in the cohort analysis and using the case–control prevalence $p_X^c$ in the case–control analysis. Under this standardization, we have

$$\left(\gamma_j^{L,c}\sqrt{p_X^c(1-p_X^c)}\right) \approx \frac{p_X^c(1-p_X^c)}{p_X(1-p_X)}\left(\gamma_j^L\sqrt{p_X(1-p_X)}\right).$$

Rearranging yields

$$\gamma_j^{L,c} \approx \frac{\sqrt{p_X^c(1-p_X^c)}}{\sqrt{p_X(1-p_X)}}\gamma_j^L \approx \frac{\sqrt{p_X^c(1-p_X^c)}}{p_X(1-p_X)}\phi(t_X)\gamma_j^\star. \tag{8}$$

#### S1.3 Approximation accuracy based on local curvature

In this section, we focus on the component of approximation error arising from the first-order Taylor expansion. A more detailed analysis of the full approximation error is provided in Section S1.4.

The accuracy of using $s_X\gamma_j$ from a binary trait to approximate the marginal association $\gamma_j^\star$ for the underlying continuous trait depends on the quality of the Taylor expansion in (2). In particular, this accuracy is determined by the magnitude of the higher-order terms omitted from the expansion. To illustrate this, we consider the second-order Taylor expansion

$$\begin{aligned} g(\mathrm{P}(X=1)) &\approx g\big(\Phi(\gamma_j^\star Z_j - t_X)\big) \\ &= (g\Phi)(-t_X) + (g\Phi)'(-t_X)\,\gamma_j^\star Z_j + \frac{1}{2}(g\Phi)''(-t_X)\,\gamma_j^{\star 2} Z_j^2 + O(\gamma_j^{\star 3}). \end{aligned} \tag{9}$$

As a result, the true regression coefficient $\gamma_j$ deviates from $s_X\gamma_j^\star$ due to the linear projection of the second-order term $\frac{1}{2}(g\Phi)''(-t_X)\,\gamma_j^{\star 2}Z_j^2$ onto $Z_j$. The sensitivity of the link function $g$ to the relative error between $\gamma_j$ and $s_X\gamma_j^\star$ is therefore governed by the ratio

$$\frac{(g\Phi)''(-t_X)}{(g\Phi)'(-t_X)},$$

which we refer to as the *relative curvature* of the link-composition function. In particular, when linear regression is used, and $g$ is the identity link, we have

$$\frac{(g\Phi)''(-t_X)}{(g\Phi)'(-t_X)} = \frac{\Phi''(-t_X)}{\Phi'(-t_X)} = t_X = \Phi^{-1}(1-p_X).$$

For logistic regression, where $g = H^{-1}$ with $H$ the logistic function, we have

$$(g\Phi)(x) = H^{-1}(\Phi(x)) = \log\left(\frac{\Phi(x)}{1-\Phi(x)}\right), \qquad (g\Phi)'(x) = \frac{\phi(x)}{\Phi(x)(1-\Phi(x))}.$$

Differentiating once more gives

$$\begin{aligned}(g\Phi)''(x) &= \frac{\phi'(x)\Phi(x)(1-\Phi(x)) - \phi(x)\big(\phi(x)(1-\Phi(x)) - \phi(x)\Phi(x)\big)}{\Phi(x)^2(1-\Phi(x))^2} \\ &= -\,\frac{\phi(x)\,[x\Phi(x)(1-\Phi(x)) + \phi(x)(1-2\Phi(x))]}{\Phi(x)^2(1-\Phi(x))^2} \\ &= -\left(x\,\frac{\phi(x)}{\Phi(x)(1-\Phi(x))} + \frac{\phi(x)^2(1-2\Phi(x))}{\Phi(x)^2(1-\Phi(x))^2}\right).\end{aligned}$$

Consequently,

$$\frac{(g\Phi)''(-t_X)}{(g\Phi)'(-t_X)} = t_X - \frac{\phi(-t_X)(1-2\Phi(-t_X))}{\Phi(-t_X)(1-\Phi(-t_X))}.$$

Since $t_X = \Phi^{-1}(1-p_X)$ and $\Phi(-t_X) = p_X$, this expression can be written as

$$\frac{(g\Phi)''(-t_X)}{(g\Phi)'(-t_X)} = \Phi^{-1}(1-p_X) - \frac{\phi\big(\Phi^{-1}(p_X)\big)(1-2p_X)}{p_X(1-p_X)}.$$

##### S1.4 Refined approximation accuracy analysis

Based on Section S1.1, our derivations of proportionality actually rely on two key approximations: (1) Assume $X_j^\star$ and $\mu_j + \epsilon_j$ roughly follow the same distribution, which is justified by the small-effect assumption $\gamma_j^\star = o(1)$; and (2) truncating the Taylor expansion at first order. Section S1.3 mainly discusses the approximation accuracy arising from the second step. In this section, we quantify the approximation error introduced by both steps under the following model:

$$\begin{aligned} B_j &\sim \text{Binomial}(2, p_j), \\ Z_j &= B_j - 2p_j, \\ X_j^\star \mid Z_j &\sim N(\gamma_j^\star Z_j, \sigma_j^2), \\ \text{Var}(X_j^\star) &= 1. \end{aligned}$$

Under this model, the marginal variance decomposition gives

$$\sigma_j^2 = \boldsymbol{E}\big[\text{Var}(X_j^\star \mid Z_j)\big] = \text{Var}(X_j^\star) - \text{Var}\big(\boldsymbol{E}[X_j^\star \mid Z_j]\big) = 1 - \gamma_j^{\star 2}\text{Var}(Z_j).$$

We now evaluate the accuracy of the approximation $s_X \approx (g\Phi)'(t_X)$. Specifically, we show that the resulting relative error in the estimated regression coefficient approximately satisfies

$$\left|\frac{\gamma_j - (g\Phi)'(t_X)\,\gamma_j^\star}{\gamma_j}\right| \le \frac{1-\sigma_j^2}{\sigma_j(1+\sigma_j)} + \left|\frac{(g\Phi)''(-t_X)}{(g\Phi)'(-t_X)}\right| M_j$$

where

$$\frac{1-\sigma_j^2}{\sigma_j(1+\sigma_j)} \approx \frac{\gamma_j^{\star 2}\,\text{Var}(Z_j)}{2}, \quad M_j = \left[\frac{|2\sigma_j - 1|}{\sigma_j^2}\left(1 + \frac{\gamma_j^{\star 2}\text{Var}(Z_j)t_X^2}{(1+\sigma_j)^2}\right)\right]|\gamma_j^\star| \approx |\gamma_j^\star|.$$

For example, with allele frequency $p_j = 0.3$, trait prevalence $p_X = 0.1$, and $\gamma_j^{\star 2}\text{Var}(Z_j) = 0.001$, the relative error is bounded above by approximately 0.015 for logistic regression and 0.063 for linear regression.

**Approximation error in Equation** (1)**.** Under these assumptions, we have

$$\begin{aligned}
\mathrm{P}(X = 1 \mid Z_j) &= \mathrm{P}(X^\star > t_X \mid Z_j) \\
&= \mathrm{P}(\gamma_j^\star Z_j + \epsilon_j > t_X \mid Z_j) \\
&= \mathrm{P}\left(\frac{-\epsilon_j}{\sigma_j} < \frac{\gamma_j^\star Z_j - t_X}{\sigma_j} \middle| Z_j\right) \\
&= \Phi\left(\frac{\gamma_j^\star}{\sigma_j} Z_j - \frac{t_X}{\sigma_j}\right),
\end{aligned} \tag{10}$$

In contrast, the approximation used earlier replaces this expression by

$$\Phi\left(\gamma_j^\star Z_j - t_X\right).$$

Thus, the approximation error in the first step arises from replacing $\gamma_j^\star/\sigma_j$ by $\gamma_j^\star$.

**Approximation in Equation (2)** Define

$$a_j := -\frac{t_X}{\sigma_j} - (-t_X) = \left(1 - \frac{1}{\sigma_j}\right) t_X$$

as the baseline adjustment. Starting from (10), we write

$$\begin{aligned}
g(\mathrm{P}(X = 1)) &= (g\Phi)\left(\frac{\gamma_j^\star}{\sigma_j} Z_j - \frac{t_X}{\sigma_j}\right) \\
&= (g\Phi)(-t_X) + (g\Phi)'(-t_X)\left(\frac{\gamma_j^\star}{\sigma_j} Z_j + a_j\right) + \frac{1}{2}(g\Phi)''(-t_X + \eta_j)\left(\frac{\gamma_j^\star}{\sigma_j} Z_j + a_j\right)^2,
\end{aligned}$$

for some $\eta_j$ satisfying $|\eta_j| \leq \left|\frac{\gamma_j^\star}{\sigma_j} Z_j + a_j\right|$. Let

$$\delta_j := \max_{b \in \{0,1,2\}} \left|\frac{\gamma_j^\star}{\sigma_j}(b - 2p_j) + a_j\right|, \quad R_j := \sup_{|\eta| \leq \delta_j} \left|(g\Phi)''(-t_X + \eta)\right|$$

Using $|Z_j| \leq 2$ and $\mathrm{E}[Z_j] = 0$, we obtain

$$\begin{aligned}
\left|\mathrm{Cov}\left(\frac{1}{2}(g\Phi)''(-t_X + \eta_j)\left(\frac{\gamma_j^\star}{\sigma_j} Z_j + a_j\right)^2, Z_j\right)\right| &= \frac{1}{2}\left|\boldsymbol{E}\left[(g\Phi)''(-t_X + \eta_j)\left(\frac{\gamma_j^\star}{\sigma_j} Z_j + a_j\right)^2 Z_j\right]\right| \\
&= \frac{1}{2}\boldsymbol{E}\left[\left|(g\Phi)''(-t_X + \eta_j) Z_j\right|\left(\frac{\gamma_j^\star}{\sigma_j} Z_j + a_j\right)^2\right] \\
&\leq R_j \boldsymbol{E}\left[\left(\frac{\gamma_j^\star}{\sigma_j} Z_j + a_j\right)^2\right] \\
&= R_j\left(\frac{\gamma_j^{\star 2}}{\sigma_j^2}\mathrm{Var}(Z_j) + a_j^2\right)
\end{aligned}$$

Consequently, the linear projection of $g(\mathrm{P}(X=1))$ onto $Z_j$ gives coefficient

$$\frac{\mathrm{Cov}(g(P(X=1)), Z_j)}{\mathrm{Var}(Z_j)} = (g\Phi)'(-t_X)\frac{\gamma_j^\star}{\sigma_j} Z_j + O_j Z_j + \tilde{\epsilon}_j,$$

where $C_j$ is a constant that does not depend on $Z_j$, $\mathrm{Cov}(Z_j, \tilde{\epsilon}_j) = 0$ and

$$|O_j| \leq \frac{1}{\mathrm{Var}(Z_j)}\left(R_j\left(\frac{\gamma_j^{\star,2}}{\sigma_j^2}\mathrm{Var}(Z_j) + a_j^2\right)\right) = R_j\left(\frac{\gamma_j^{\star 2}}{\sigma_j^2} + \frac{a_j^2}{\mathrm{Var}(Z_j)}\right)$$

**Relative approximation error.** Comparing with the first-order approximation

$$g(\mathrm{P}(X=1)) \approx (g\Phi)(-t_X) + (g\Phi)'(-t_X)\,\gamma_j^\star Z_j,$$

the relative error in the regression coefficient satisfies

$$\begin{aligned}\left|\frac{(g\Phi)'(-t_X)\frac{\gamma_j^\star}{\sigma_j} + O_j - (g\Phi)'(-t_X)\gamma_j^\star}{(g\Phi)'(-t_X)\frac{\gamma_j^\star}{\sigma_j} + O_j}\right| &\leq \left|\frac{(g\Phi)'(-t_X)\frac{\gamma_j^\star}{\sigma_j} + \frac{O_j}{\sigma_j} - (g\Phi)'(-t_X)\gamma_j^\star - O_j}{(g\Phi)'(-t_X)\frac{\gamma_j^\star}{\sigma_j} + O_j}\right| \\ &\quad + \left|\frac{(2-\frac{1}{\sigma_j})O_j}{(g\Phi)'(-t_X)\frac{\gamma_j^\star}{\sigma_j} + O_j}\right| \\ &= \left(\frac{1}{\sigma_j} - 1\right) + \left|\frac{(2\sigma_j - 1)O_j}{(g\Phi)'(-t_X)\gamma_j^\star + \sigma_j O_j}\right|\end{aligned}$$

The first term has closed from

$$\left(\frac{1}{\sigma_j} - 1\right) = \frac{(1-\sigma_j)(1+\sigma_j)}{\sigma_j(1+\sigma_j)} = \frac{\gamma_j^{\star 2}\mathrm{Var}(Z_j)}{\sigma_j(1+\sigma_j)} = \frac{\gamma_j^{\star 2}\mathrm{Var}(Z_j)}{\sqrt{1-\gamma_j^{\star 2}\mathrm{Var}(Z_j)} + (1-\gamma_j^{\star 2}\mathrm{Var}(Z_j))}$$

where $\gamma_j^{\star 2}\mathrm{Var}(Z_j)$ represents the proportion of variance of $X^\star$ explained by SNP $Z_j$. This quantity, depending only on the variance explained by the SNP $j$ is less than 0.0541 when $\gamma_j^{\star 2}\mathrm{Var}(Z_j) \leq 0.1$, and less than 0.026 when $\gamma_j^{\star 2}\mathrm{Var}(Z_j) \leq 0.05$.

For the second term, first note

$$\begin{aligned}|O_j| &\leq R_j\left(\frac{\gamma_j^{\star 2}}{\sigma_j^2} + \frac{a_j^2}{\mathrm{Var}(Z_j)}\right) \\ &= R_j\left(\frac{\gamma_j^{\star 2}}{\sigma_j^2} + \left(\frac{1}{\sigma_j} - 1\right)^2 \frac{t_X^2}{\mathrm{Var}(Z_j)}\right) \\ &= R_j\left(\frac{\gamma_j^{\star 2}}{\sigma_j^2} + \frac{\gamma_j^{\star 4}\mathrm{Var}(Z_j)^2}{\sigma_j^2(1+\sigma_j)^2}\frac{t_X^2}{\mathrm{Var}(Z_j)}\right) \\ &= R_j\frac{\gamma_j^{\star 2}}{\sigma_j^2}\left(1 + \frac{\gamma_j^{\star 2}\mathrm{Var}(Z_j)t_X^2}{(1+\sigma_j)^2}\right)\end{aligned}$$

Therefore

$$\left|\frac{(2\sigma_j-1)O_j}{(g\Phi)'(-t_X)\gamma_j^\star+\sigma_jO_j}\right| \le \left|\frac{2\sigma_j-1}{(g\Phi)'(-t_X)\gamma_j^\star+\sigma_jO_j}\right| \frac{R_j\gamma_j^{\star 2}}{\sigma_j^2}\left(1+\frac{\gamma_j^{\star 2}\mathrm{Var}(Z_j)t_X^2}{(1+\sigma_j)^2}\right)$$
$$= \left|\frac{R_j}{(g\Phi)'(-t_X)+\sigma_jO_j/\gamma_j^\star}\right| \frac{|2\sigma_j-1|}{\sigma_j^2}\left(1+\frac{\gamma_j^{\star 2}\mathrm{Var}(Z_j)t_X^2}{(1+\sigma_j)^2}\right)|\gamma_j^\star|$$

Note the only component depending on the link function is

$$\frac{R_j}{(g\Phi)'(-t_X)+\sigma_jO_j/\gamma_j^\star} = \sup_{|\eta|\le\delta_j}\frac{\left|(g\Phi)''(-t_X+\eta)\right|}{(g\Phi)'(-t_X)+\sigma_jO_j/\gamma_j^\star}, \quad \delta_j = \max_{b\in\{0,1,2\}}\left|\frac{\gamma_j^\star}{\sigma_j}(b-2p_j)+a_j\right|.$$

Since $\delta_j = O(\gamma_j^\star)$ and $O_j/\gamma_j^\star = O(\gamma_j^\star)$ are small, this quantity can be well-approximated by evaluating the ratio at the expansion point,

$$\frac{(g\Phi)''(-t_X)}{(g\Phi)'(-t_X)},$$

which is the relative curvature defined in Section S1.3.

### S2 Interpretation under nonlinear causal effect of $X^\star$ on $Y^\star$

In Section 3.4 of the maintext, we began by noting that, under standard MR assumptions imposed on the latent liabilities, the genetic associations satisfy

$$\Gamma_j^\star = \beta\gamma_j^\star + \alpha_j^\star,$$

We then showed that, on the observed scale, this relationship implies $\Gamma_j \approx \beta\frac{s_Y}{s_X}\gamma_j + s_Y\alpha_j^\star$. This result on summary statistics relies on the assumption that the latent liabilities follow the linear structural model

$$Y^\star = \beta X^\star + f_0(\boldsymbol{Z},\boldsymbol{U},\boldsymbol{E}_Y),$$

where $f_0$ is a differentiable function, $\boldsymbol{Z} = (Z_j)_{j=1}^p$ denotes the set of genetic variants, $\boldsymbol{U}$ represents unmeasured confounders, and $\boldsymbol{E}_Y$ denotes exogenous noise. In this section, we examine several ways in which the linear causal effect of $X^\star$ on $Y^\star$ may be violated and discuss the implications for our results.

#### S2.1 Nonlinear smooth relationships between $X^\star$ and $Y^\star$

In Supplement Section S1.1.3 of our earlier work [6], we considered the interpretation of MR when the latent exposure and outcome follow a nonlinear structural model of the form

$$Y^\star = f_0(X^\star,\boldsymbol{Z},\boldsymbol{U},\boldsymbol{E}_Y),$$

where $f_0$ is a differentiable function. In this setting, they show that the marginal associations between SNP $j$ and the latent traits satisfy

$$\Gamma_j^\star \approx \beta^\star \gamma_j^\star + \alpha_j^\star, \qquad \beta^\star = \boldsymbol{E}\left[\frac{\partial}{\partial x} f_0(X^\star, \boldsymbol{Z}, \boldsymbol{U}, \boldsymbol{E}_Y)\right],$$

where $\beta^\star$ is the *average derivative effect* (ADE [3]) and $\alpha_j^\star$ is a residual term defined in [6].

This analysis does not directly extend to the observed binary outcome $Y = \mathbf{1}\{Y^\star > c\}$, since the indicator transformation is not differentiable and therefore falls outside the scope of the ADE framework. Nevertheless, by applying the proportionality arguments developed in this paper, we can still obtain an approximate relationship for the observed-scale associations:

$$\Gamma_j \approx \frac{s_Y}{s_X} \beta^\star \gamma_j + s_Y \alpha_j^\star,$$

indicating that the same linear form is preserved up to the appropriate scaling factors, even in the presence of nonlinear latent effects. In this case, the MR coefficient should be interpreted as a scaled version of the ADE.

##### S2.2 Stepwise causal effects of $X^\star$ on $Y^\star$

As discussed in [4], the effect of a latent liability on an outcome may operate not only directly through the continuous liability itself, but also indirectly through a categorical threshold. For example, individuals born with an orofacial cleft typically undergo corrective surgery, whereas individuals who do not develop an orofacial cleft do not, regardless of their underlying liability. In this setting, the latent liability influences the outcome primarily through the binary indicator of whether the threshold is crossed. Similarly, liability to educational attainment may affect downstream outcomes, such as the number of books read, partly through the attainment of formal educational qualifications.

To capture such stepwise mechanisms, consider the generating model

$$Y^\star = \beta_1 X^\star + \beta_2 X + f_0(\boldsymbol{Z}, \boldsymbol{U}, \boldsymbol{E}_Y) = \beta_1 X^\star + \beta_2\, 1(X^\star > t_X) + f_0(\boldsymbol{Z}, \boldsymbol{U}, \boldsymbol{E}_Y),$$

so that the relationship between $Y^\star$ and $X^\star$ is nonlinear and non-differentiable. Under this specification, the marginal association between SNP $j$ and the latent outcome satisfies

$$\Gamma_j^\star = \beta_1 \gamma_j^\star + \beta_2 \gamma_j + \alpha_j^\star,$$

where $\alpha_j^\star$ represents the coefficient obtained by projecting $f_0(\boldsymbol{Z}, \boldsymbol{U}, \boldsymbol{E}_Y)$ linearly onto $Z_j$. Consequently, on the observed scale we obtain

$$\Gamma_j \approx s_Y \left(\frac{1}{s_X} \beta_1 + \beta_2\right) \gamma_j + s_Y\, \alpha_j^\star,$$

showing that the MR coefficient corresponds to a scaled weighted combination of $\beta_1$ and $\beta_2$, where the weights depend on the prevalence of the binary traits.

#### S3 Extension of the scaling framework

##### S3.1 GWAS summary statistics after adjusting for additional covariates

In practice, GWAS summary statistics are often obtained after adjusting for additional environmental covariates $\boldsymbol{C}$, such as age, sex, and leading principal components capturing population structure. Under standard Mendelian randomization assumptions, genetic variants $\boldsymbol{Z}$ are unconfounded and therefore independent of these environmental covariates. As a result, our earlier argument—that regression coefficients from logistic or linear models fitted to binary traits are approximately proportional to those from linear regression on the underlying continuous traits—continues to hold under covariate adjustment.

Under the assumption that $\boldsymbol{C} \perp\!\!\!\perp Z_j$, coefficients from linear regression are invariant to adjustment: both the coefficient $\gamma_j^\star$ for the continuous trait $X^\star$ and the coefficient $\gamma_j^L$ for the binary trait $X$ remain unchanged whether or not $\boldsymbol{C}$ is included in the model. In contrast, logistic regression coefficients are affected by covariate adjustment due to the well-known non-collapsibility of odds ratios [2].

To illustrate this effect, consider the logistic regression model adjusted for $\boldsymbol{C}$,

$$\mathrm{P}(X = 1 \mid Z_j, \boldsymbol{C}) = H\Big(\gamma_{0j,C} + \gamma_{j,C}^H Z_j + \boldsymbol{C}^\top \boldsymbol{\beta}_j\Big).$$

Since the effect of any single SNP is typically small, the coefficients $\boldsymbol{\beta}_j$ associated with the environmental covariates are expected to be similar across $j$. Moreover, when the prevalence $\mathrm{P}(X = 1)$ is not extreme and $\boldsymbol{C}$ is approximately normally distributed, the approximation $H(x) \approx \Phi(x/1.7)$ can be applied. Following arguments similar to those in [7], this yields

$$\mathrm{P}(X = 1 \mid Z_j) \approx H\left(\frac{\gamma_{0j,C} + \gamma_{j,C}^H Z_j}{\sqrt{1 + \mathrm{Var}(\boldsymbol{C}^\top \boldsymbol{\beta}_j)/1.7^2}}\right).$$

Given that $\boldsymbol{\beta}_j$ is similar across $j$, we denote the common value by $\boldsymbol{\beta}$. Under this approximation, the adjusted logistic regression coefficient $\gamma_{j,C}^H$ is approximately proportional to the unadjusted coefficient $\gamma_j^H$, and hence also proportional to the regression coefficient for the underlying continuous trait. The proportionality factor depends on the contribution of $\boldsymbol{C}$ to the binary trait $X$:

$$\gamma_{j,C}^H \approx \sqrt{1 + \mathrm{Var}(\boldsymbol{C}^\top \boldsymbol{\beta})/1.7^2}\,\gamma_j^H \approx \sqrt{1 + \mathrm{Var}(\boldsymbol{C}^\top \boldsymbol{\beta})/1.7^2}\,\frac{\phi(t_X)}{p_X(1 - p_X)}\gamma_j^\star \tag{11}$$

##### S3.2 Summary statistics on ordered categorical traits

In GWAS, many phenotypes are ordinal rather than binary. We show that, under standard assumptions, regression coefficients for ordered categorical traits remain approximately proportional to the marginal effect $\gamma_j^\star$ of the underlying continuous trait.

As in the binary-trait setting, we model an ordered categorical trait as a discretization of an underlying continuous liability $X^\star$, which we assume follows a standard normal distribution,

$X^\star \sim N(0,1)$. Specifically, for an ordered categorical variable $X$ with $K$ levels, we assume

$$X = k \quad \text{if} \quad X^\star \in (t_{k-1}, t_k],$$

where $k \in \{1, \dots, K\}$ and

$$-\infty = t_0 < t_1 < \cdots < t_K = \infty.$$

Assuming that the marginal SNP effect is small, $\gamma_j^\star = o(1)$, we have

$$\begin{aligned} \mathrm{P}(X \le k \mid Z_j) &= \mathrm{P}(X^\star \le t_k \mid Z_j) \\ &= \mathrm{P}(\mu_j + \gamma_j^\star Z_j + \epsilon_j \le t_k \mid Z_j) \\ &\approx \Phi(t_k - \gamma_j^\star Z_j), \end{aligned} \tag{12}$$

where the approximation follows from the small magnitude of $\gamma_j^\star$.

**Ordinal regression.** In cumulative link ordinal regression models, we assume

$$H^{-1}(\mathrm{P}(X \le k \mid Z_j)) = \gamma_{0j,k} - \gamma_j^H Z_j,$$

where $H(\cdot)$ denotes the logistic link function. When the probabilities $\mathrm{P}(X = 1)$ and $\mathrm{P}(X = K)$ are not extreme, the approximation $H(x) \approx \Phi(x/1.7)$ can be applied. This yields

$$\mathrm{P}(X \le k \mid Z_j) = H\big(\gamma_{0j,k} - \gamma_j^H Z_j\big) \approx \Phi\left(\frac{\gamma_{0j,k} - \gamma_j^H Z_j}{1.7}\right).$$

Comparing this expression with (12) implies

$$\gamma_j^H \approx s_X \gamma_j^\star, \qquad s_X = 1.7. \tag{13}$$

**Linear regression.** Alternatively, suppose the ordinal outcome $X$ is analyzed using linear regression to obtain GWAS summary statistics. The fitted model is

$$\boldsymbol{E}[X \mid Z_j] = \gamma_{0j} + \gamma_j^L Z_j.$$

Using the tail-sum representation of expectations,

$$\begin{aligned}
\boldsymbol{E}[X \mid Z_j] &= \sum_{k=0}^{K-1} \mathrm{P}(X > k \mid Z_j) \\
&= K - \sum_{k=0}^{K-1} \mathrm{P}(X \leq k \mid Z_j) \\
&\approx K - \sum_{k=0}^{K-1} \Phi(t_k - \gamma_j^\star Z_j) \\
&\approx K - \sum_{k=0}^{K-1} \left[\Phi(t_k) - \phi(t_k)\, \gamma_j^\star Z_j\right] \\
&= \left(K - \sum_{k=1}^{K-1} \Phi(t_k)\right) + \left(\sum_{k=1}^{K-1} \phi(t_k)\right) \gamma_j^\star Z_j.
\end{aligned}$$

This expansion implies that

$$\gamma_j^L \approx s_X \gamma_j^\star, \qquad s_X = \sum_{k=1}^{K-1} \phi(t_k), \tag{14}$$

where the scaling factor $s_X$ depends only on the cutpoints $\{t_k\}$ and is therefore constant across SNPs.

##### S3.3 GWAS summary statistics for binary traits under liability distribution

So far, we have assumed that the latent trait $X^\star$ follows a standard normal distribution. Here we emphasize that the same proportionality argument holds more generally, without requiring Gaussianity of the latent variable.

Specifically, let $F$ denote the cumulative distribution function (CDF) of $-X^\star$. Under the small-effect assumption $\gamma_j^\star = o(1)$, the Taylor expansion arguments used previously remain valid for any smooth distribution $F$. Consequently, the relationship between the observed binary-trait regression coefficient and the latent continuous effect can be written in the general form

$$\gamma_j \approx s_X \gamma_j^\star, \quad s_X = (gF)'\big(F^{-1}(p_X)\big), \tag{15}$$

where $g(\cdot)$ is the link function used in the binary regression model and $p_X = \mathrm{P}(X = 1)$ is the trait prevalence.

As before, the accuracy of this first-order approximation depends on the magnitude of the higher-order terms. The relevant measure of sensitivity is characterized by the curvature of the composite function $gF$, given by

$$\frac{(gF)''(-t_X)}{(gF)'(-t_X)} = \frac{(gF)''(F^{-1}(p_X))}{(gF)'(F^{-1}(p_X))}$$

#### S4 Simulation details

Here we provide more details of our simulation setup. First we generate the effect size $\gamma_j^*$ of 100 independent SNPs $j$ on the trait $X^*$ from standard Gaussian distribution

$$\gamma_j \sim \mathcal{N}(0,1), \quad j = 1, 2, \cdots, 100,$$

The EAFs $(p_1, \cdots, p_{100})$ of the SNPs are randomly sampled from the empirical EAFs in the UK Biobank.

Then we conduct experiments for 100 times with these fixed effect sizes and EAFs. At each replication, we generate a population of $n = 400K$ individuals. Specifically, the SNPs $Z_{ij}$, the continuous traits $X_i^*$ and the binary traits $X_i$ are generated following:

$$\begin{aligned}
Z_{ij} &\sim \text{Bernoulli}(p_j), \quad i = 1, 2, \cdots, 4 \cdot 10^5, \\
\epsilon_i &\sim \mathcal{N}\left(0, \text{Var}\left(\sum_{j=1}^{100} Z_{ij}\gamma_j\right)\right), \\
\tilde{X}_i^\star &= \sum_{j=1}^{100} Z_{ij}\gamma_j + \epsilon_i, \\
X_i^\star &= \left(\tilde{X}_i^\star - \boldsymbol{E}[\tilde{X}_i^\star]\right) / \sqrt{\text{Var}(\tilde{X}_i^\star)}, \\
X_i &= \mathbf{1}\{X_i^\star \geq \text{quantile}(X_i^\star, c)\}, \quad c = 0.5, 0.8, 0.95, 0.99, 0.999.
\end{aligned}$$

Here the variance of $\epsilon_i$ is designed in the way such that these 100 SNPs in total explain approximately 60% of the variability in $X$. The threshold $c$ is chosen such that $\text{P}(X = 1) = 0.5, 0.2, 0.05, 0.01, 0.001$, respectively. Here $\tilde{X}_i^\star$ represents the unscaled trait and $X_i^\star$ represents the standardized trait. And we let

$$\gamma_j^\star = \gamma_j / \sqrt{\text{Var}(\tilde{X}_i^\star)}$$

be the true effect sizes of $Z_j$ on the standardized $X^\star$. Since the EAFs can be relatively low for some SNPs, quasi-complete separation could occur in these 100 replications for logistic regression estimates. Here we just remove the logistic regression estimates for such SNPs if such computational errors occur.

For the evaluation of MR results, we generate both the exposure and outcome traits in a similar approach as discussed above. Specifically, we generate the continuous exposure trait $X^\star$ as above and set the binary trait $X$ to have a prevalence $c$:

$$X_i = \mathbf{1}\{X_i^\star \geq \text{quantile}(X_i^\star, c)\}, \quad c = 0.5, 0.95, 0.99$$

for individual $i$. Then we generate outcome traits $Y$ and $Y^\star$ given the continuous exposure trait

$X^\star$ and the pleiotropic effects $\alpha$:

$$
\begin{aligned}
\alpha_j &\sim \mathcal{N}(0,\sigma^2), \quad \sigma = 0.008 \text{ or } 0.08, \quad j = 1, 2, \cdots, 100,\\
\epsilon_i^Y &\sim \mathcal{N}(0, 0.7),\\
\tilde{Y}_i^\star &= X_i^\star + \sum_{j=1}^{100} Z_{ij}\alpha_j + \epsilon_i^Y,\\
Y_i^\star &= \left(\tilde{Y}_i^\star - \boldsymbol{E}[\tilde{Y}_i^\star]\right)/\sqrt{\mathrm{Var}(\tilde{Y}_i^\star)},\\
Y_i &= \mathbf{1}\{Y_i^\star \geq \mathrm{quantile}(Y_i^\star, c)\}, \quad c = 0.5, 0.95, 0.99\\
\beta &= 1/\sqrt{\mathrm{Var}(\tilde{Y}_i^\star)}.
\end{aligned}
$$

Here $\sigma = 0.08$ corresponds to the medium pleiotropy setting and $\sigma = 0.008$ corresponds to the weak pleiotropy setting. The parameter $\beta$ represents the causal effect of a one-unit increase in the latent exposure $X^\star$ on a one-unit increase in the latent outcome $Y^\star$. The Inverse-variance weighted (IVW) analyses were performed using the MVMR package (`https://github.com/WSpiller/MVMR` ), and GRAPPLE analyses were conducted using the *grappleRobustEst* function from the GRAPPLE package (`https://github.com/jingshuw/GRAPPLE`). Prior to MR analysis, we constructed Z-scores using $\gamma_j^\star/\mathrm{SE}(\hat{\gamma}_j{}^H)$ from the logistic regression, and selected significant SNPs with $p$-values below $10^{-8}$.

#### S5 Supplementary Figures

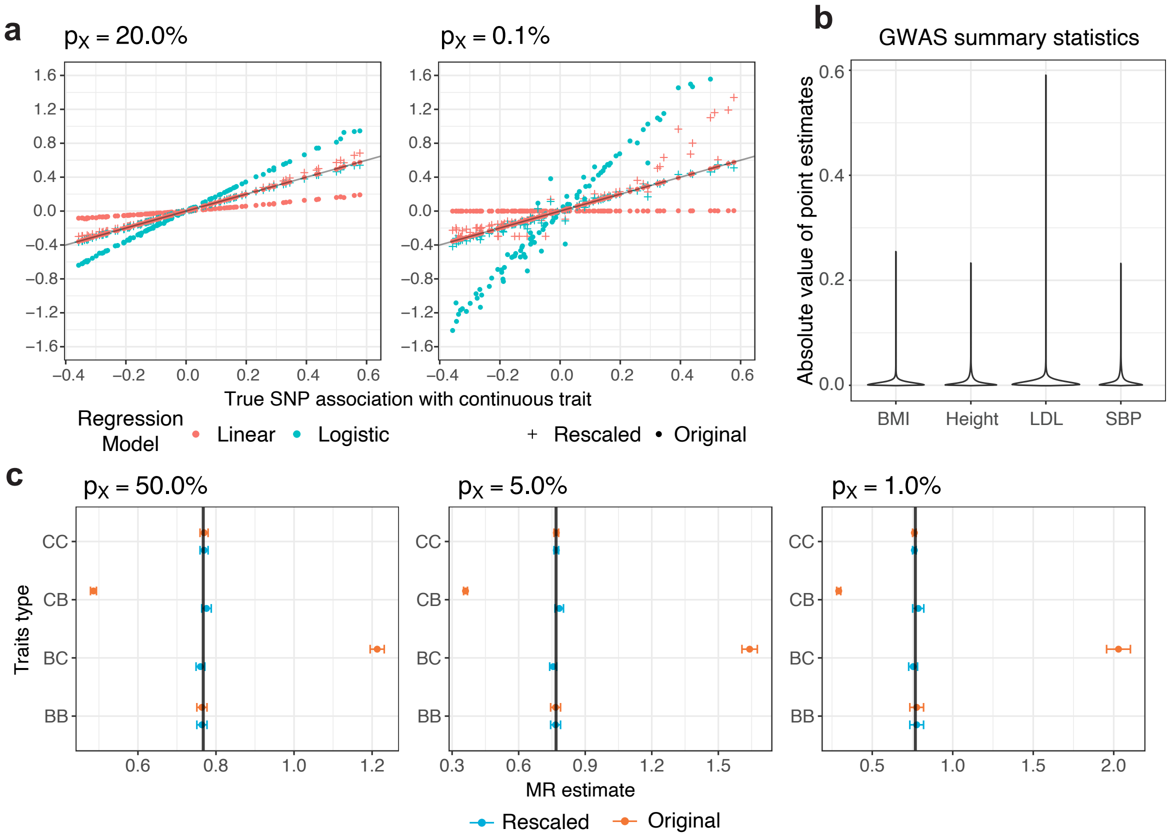


Figure S1: a) True per-SNP associations on the underlying continuous trait versus associations estimated from the observed binary trait using logistic or linear regression, shown before and after rescaling by the factor $s_X$, for prevalences $p_X = 20\%$ and $0.1\%$. Rescaled estimates are obtained by dividing the binary-trait regression coefficients by $s_X$. Points represent median estimates across 100 simulation replicates. The black line indicates $y = x$. b) Range of observed summary statistics on common continuous traits. c) MR 95% confidence intervals computed using GRAPPLE under small pleiotropy, shown before and after rescaling, for four exposure-outcome combinations: continuous-continuous (CC), continuous-binary (CB), binary-continuous (BC), and binary-binary (BB). When traits are binary, exposure and outcome share the same prevalence. Results are shown for one representative simulation replicate. The true causal effect is approximately 0.75 (black vertical line).
